# CCAAT-enhancer binding protein delta functions as a tumor suppressor gene in acute myeloid leukemia

**DOI:** 10.1101/2025.08.25.670309

**Authors:** Subhash C. Prajapati, Cem Meydan, Yaseswini Neelamraju, Zhenjia Wang, Hao Fan, Nicholas Dunham, Richard Dillon, Jorge A Gandara, Tak Lee, Caroline Sheridan, Paul Zumbo, Michael W. Becker, Lars Bullinger, Martin P Carroll, Richard J D’Andrea, Ross L Levine, Christopher Mason, Ari M Melnick, Chongzhi Zang, Stefan Bekiranov, Francine E. Garrett-Bakelman

## Abstract

There is a continued need for identification of novel disease drivers of acute myeloid leukemia as many patients experience relapse and have poor clinical outcomes. Analyses from our study and publicly available datasets predicted CEBPD as a novel tumor suppressor gene in acute myeloid leukemia. Consistent with the analyses, CEBPD knockdown experiments showed activation of MAPK signaling with concomitant increase in cell growth rate, while upregulation experiments suggested induction of myeloid differentiation marker CD14 expression in AML cell lines OCI-AML2 and OCI-AML5. Consistent with a previous report, our genomics analyses and azacytidine treatment experiments suggested a role for DNA methylation in downregulation of CEBPD expression during AML pathogenesis. Altogether, our results provide experimental evidence for a tumor suppressor function of CEBPD in AML.

---

Acute myeloid leukemia (AML) remains a clinical challenge as many patients experience disease relapse and have poor clinical outcomes (1). Despite discoveries of disease pathogenesis(2), our understanding of disease biology remains incomplete. There is continued need for identification of new disease drivers to improve therapeutics strategies and better manage this malignancy.

Aberrant transcriptional patterns are associated with AML pathogenesis and disease prognosis (3). Targeting of their function has shown promising results in vitro in further improving treatment options for the disease (4). To identify novel aberrantly expressed genes associated with disease pathogenesis, we performed differential gene expression analyses on a previously described study dataset (5, 6). The dataset comprised of two cohorts (I: n = 29 and II: n = 30, Supplementary table 1). Each cohort comprised matched diagnosis-relapse pairs along with corresponding healthy specimens from CD34+ normal bone marrow cells. Differential gene expression analyses after correction for batch effects (Supplementary Fig. 1A) on both the cohorts revealed concordant significant differential expression of several genes upon disease relapse (Fig. 1A, Supplementary table 2). The differentially expressed genes (DEGs) set included several genes that were previously associated with AML pathogenesis (Fig. 1A). To identify novel AML drivers, we assessed genes in the DEGs set based on the following criteria (1) concordant differential expression in high proportion of study patients upon relapse (Fig. 1A, Supplementary Fig. 1B, Supplementary table 2), (2) differential expression in independent publicly available datasets, (3) no or limited previous information on their involvement in AML pathogenesis, and (4) suggested tumor driver function in cancers other than AML. We identified XRCC2, STAT4, RBM47, and CEBPD as novel AML drivers and prioritized their further validation in AML pathogenesis. CEBPD was significantly downregulated in a large proportion (45%) of relapse patients (Supplementary Fig. 1B, Supplementary table 2). Consistent with this finding, analyses of independent publicly available datasets, validated downregulation of CEBPD expression upon AML relapse (Supplementary Fig. 1C-E). We assessed potential transcriptional regulators of the DEGs in our study dataset cohorts using binding analysis for regulation of transcription (7). The analysis predicted CEBPD as one of the transcriptional regulators for a subset of DEGs which were downregulated between normal CD34+ cells and AML diagnosis patients as well as between diagnosis and relapsed patients (Fig. 1B & Supplementary table 3-5). Moreover, survival analyses on the TCGA and TARGET datasets suggested that low CEBPD expression was associated with poor survival of AML patients (Fig. 1C, Supplementary Fig. 1F). These results altogether suggested that CEBPD may function as an AML tumor suppressor gene.

**Figure 1.**
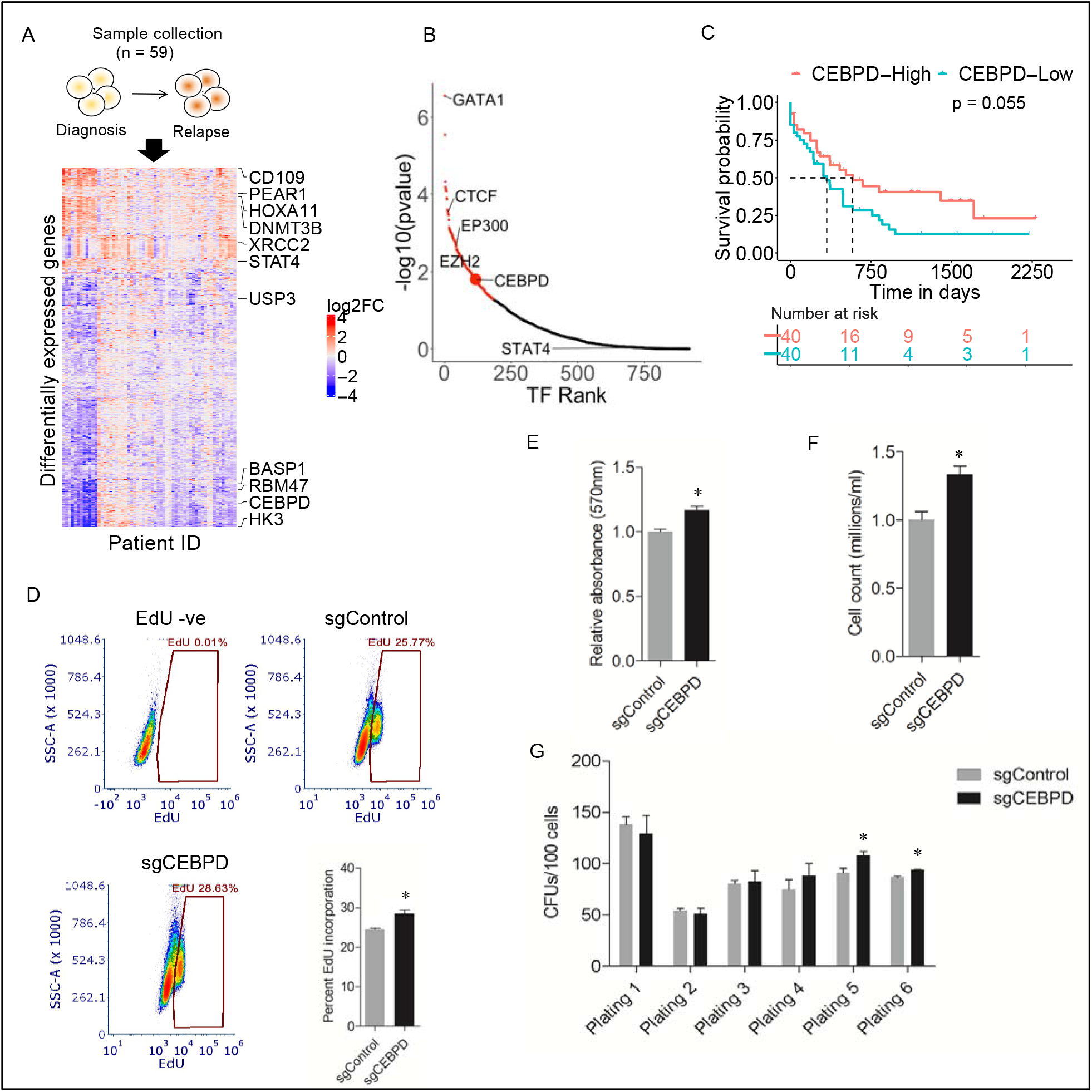
CEBPD functions as a tumor suppressor gene in AML. (A) Study specimen collection schematic and heatmap showing DEGs between patient-matched diagnosis and relapse specimens. Presented genes (n = 944) were differentially expressed in at least 25% of patients at a threshold of adjusted p-value < 0.01 and log2 fold change > |1.2|. Candidate genes together with previously implicated drivers of AML are annotated with their respective gene symbols. (B) Ranked results of predicted transcriptional regulators (TFs) from binding analysis for regulation of transcription of significantly downregulated genes in both the study cohorts. TF Rank is an arbitrary rank of the TFs. Statistically significant TFs shown in red (p-value < 0.05). (C) Kaplan Meir survival analysis results for association between CEBPD expression and overall survival in the TCGA dataset. The top and bottom quartiles of patients based on CEBPD expression were classified as CEBPD-High and CEBPD-Low, respectively. Log-rank test was used to determine the statistical significance of association. (D-H) Representative EdU incorporation assay results (D), colorimetric cell proliferation and survival analysis (E), flow cytometry-based live cell counts analysis (F), and colony forming unit (CFU) assay results (G) for CRISPR-mediated CEBPD knockdown (sgCEBPD) compared to control (sgControl) OCI-AML5 cells. Each experimental group represents an average measurement from 4 independent CEBPD-targeting and control sgRNAs. Each experimental group in the CFU assay results represents an average colony count from 3 independent CEBPD-targeting and control sgRNAs. The student’s t-test was used to determine statistical significance between the two groups.

CEBPD is a bZIP transcription factor belonging to the CCAAT enhancer binding protein (CEBP) family. Aberrant CEBPD expression has been associated with solid cancer pathogenesis as an oncogene or a tumor suppressor (8). In addition, ectopic CEBPD expression has been shown to suppress differentiation associated growth of mouse primary hematopoietic progenitor and transformed 32D-FLT3-ITD cells (9). However, the role of CEBPD in human AML pathogenesis remains poorly understood.

To determine if CEBPD can function as a tumor suppressor in human AML disease, we performed loss of function experiments in AML cell lines in which we assessed CEBPD expression (Supplementary Fig. 2A). Knockdown of CEBPD expression in the cell line OCI-AML5 using a CRISPR-mediated approach (Supplementary Fig. 2B-C) associated with enhanced cell proliferation and survival assessed using EdU incorporation (Fig. 1D) and MTS assays (Fig. 1E) as well as and flow cytometric cell counting (Fig. 1F). This finding was validated in the OCI-AML2 and OCI-AML5 cell lines using an orthogonal shRNA-mediated CEBPD knockdown approach (Supplementary Fig. 3A-D). CEBPD knockdown was also associated with enhanced self-renewal capacity in a colony forming unit assay in OCI-AML5 cells (Fig. 1G, Supplementary Fig. 3E). Furthermore, CEBPD knockdown associated with a trend of reduced apoptosis compared to controls assessed using Annexin V apoptosis assay (Supplementary Fig. 3F). We performed a CEBPD overexpression experiment in these cell models and assessed for changes in differentiation markers. Upregulation of CEBPD expression associated with higher expression of the mature myeloid marker CD14 in OCI-AML2 (Supplementary Fig. 3G-H) and OCI-AML5 cells (Supplementary Fig. 3I-J). Collectively, these results were consistent with the predicted tumor suppressor function of CEBPD.

To investigate the underlying molecular mechanism by which CEBPD knockdown promotes AML cell growth, we performed transcriptome-wide differential gene expression analyses in CRISPR-mediated CEBPD knockdown OCI-AML5 cells (Supplementary Fig. 4A, Supplementary table 6). Consistent with an enhanced cell growth rate, gene set enrichment analyses (GSEA) suggested activation of the MAPK pathway (Fig. 2A), a signaling cascade that plays a vital role in growth and survival of cancer cells (10). Consistent with the prediction, we detected elevated phosphorylation of MEK1/2 and ERK1/2 in CRISPR experiments (Fig. 2B) which confirmed activation of MAPK signaling in CEBPD knockdown cells. MAPK signaling has been associated with upregulation of downstream transducers cyclin D1 and TNF_α_, which have been associated with AML pathogenesis (11). Consistent with these reports, we observed upregulation of cyclin D1 and TNF_α_ in CEBPD knockdown cells in both CRISPR (Fig. 2C-E) and shRNA experiments (Supplementary Fig. 4B-E). Conversely, CEBPD expression has also been associated with inflammatory cytokines’ expression (12). However, these effects appear to be cell type and context dependent (12). Our observation of activation of inflammatory signaling upon loss of CEBPD expression is consistent with a previous study which showed upregulation of proinflammatory cytokines including TNF⍰ in the intestinal tissue of CEBPD null mice (13). Finally, consistent with our WB results, CEBPD knockdown associated gene expression pattern associated with sensitivity to MAPK inhibitors including trametinib and selumetinib (Fig. 2F). Collectively, our results suggest that activated MAPK signaling upon CEBPD knockdown promotes AML cell growth.

**Figure 2.**
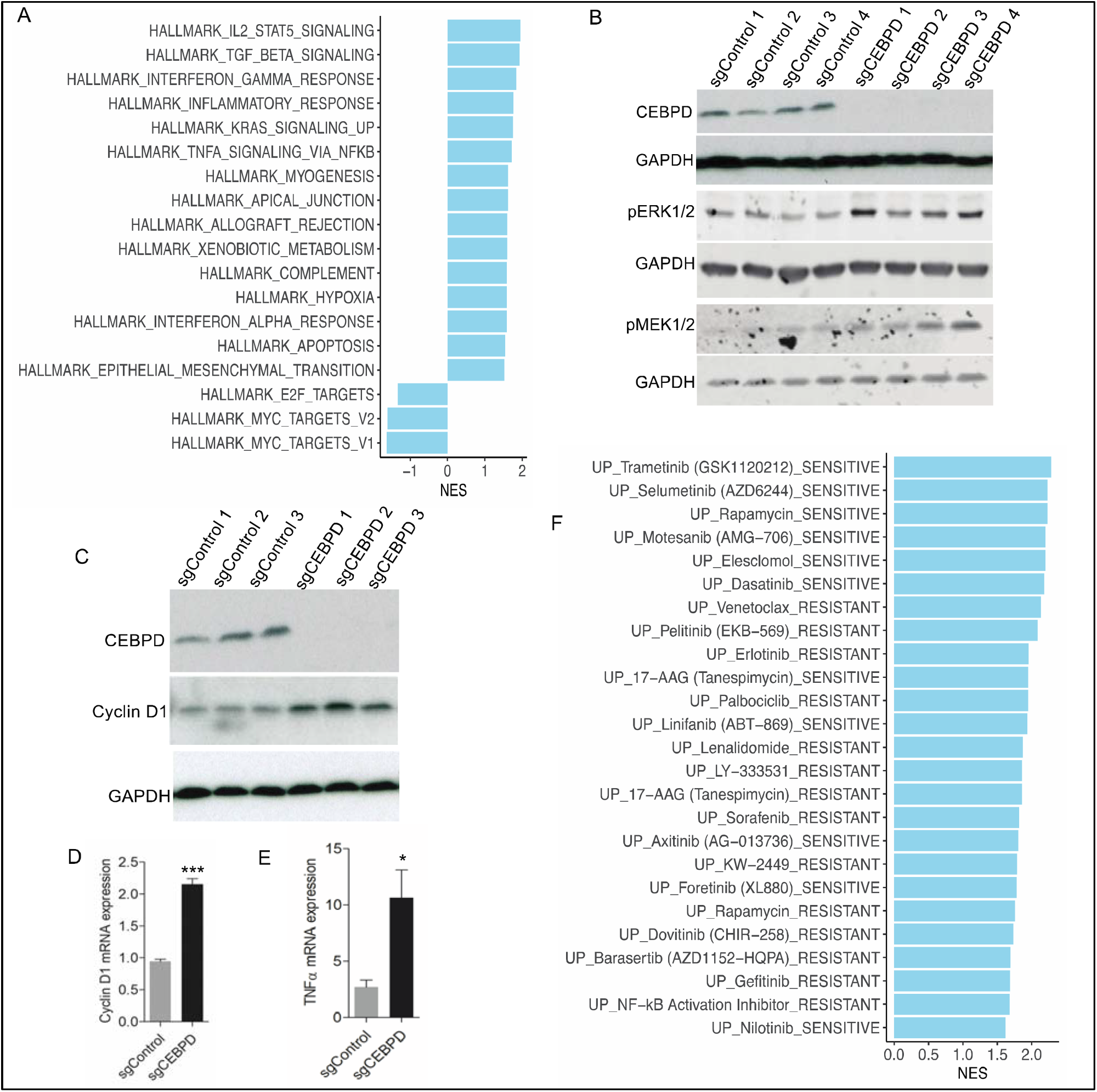
Loss of CEBPD expression is associated with activation of MAPK signaling in OCI-AML5 cells. (A) Results of fast GSEA analysis performed on the gene expression patterns in CRISPR-mediated CEBPD knockdown cells for prediction of enriched key regulatory pathways. Significantly enriched pathways were selected at a statistical threshold padj < 0.05. NES = normalized enrichment score. (B) Western blot (WB) results for pERK1/2 and pMEK1/2 levels in CEBPD knockdown (sgCEBPD) compared to control (sgControl) cells. (C) WB results for cyclin D1 expression in sgCEBPD compared to sgControl cells. (D-E) Results of qPCR analysis for cyclin D1 (D) and TNF_α_ (E) mRNA expression in sgCEBPD compared to sgControl cells, respectively. GAPDH served as a loading control and reference gene for reading normalization in all WB and qPCR analyses, respectively. (F) Results of fast GSEA analysis performed on the gene expression patterns in CRISPR-mediated CEBPD knockdown cells using treatment sensitivity gene sets from Beat AML study for prediction of sensitivity to MAPK inhibitors. Significantly enriched inhibitors were selected at a statistical threshold padj < 0.05. NES = normalized enrichment score.

We next aimed to determine the mechanisms by which CEBPD could be downregulated in during AML disease progression. Genetic alterations modify genes’ functions and are associated with pathogenesis of cancers, including AML (2). We first assessed the association between genetic alterations and CEBPD expression in AML. Copy number aberrations (CNA) analyses suggested no significant change in the CEBPD gene locus. The analyses identified one relapsed specimen in the study dataset (14) and one specimen in the TCGA dataset with gene amplification (Supplementary Fig. 5A and B). No CNA events were detected in the CEBPD gene locus in the TARGET and the Beat AML dataset specimens (Supplementary Fig. 5B). Additionally, consistent with low mutation rates in solid cancers (8), no somatic mutation was detected in the CEBPD gene in the study dataset (14) and the publicly available datasets (Supplementary Fig. 5C). These results suggest that downregulation of CEBPD expression was not associated with genetic alterations in AML. Previous reports suggest a role of DNA methylation in downregulation of CEBPD gene expression in cancers (15), including AML (9).

Although in most patients, the promoter was hypomethylated, consistent with prior reports, our analyses found a trend towards negative association between the proximal promoter methylation and CEBPD expression (Supplementary Fig. 6A-C) in the publicly available and the study datasets (Supplementary Fig. 6D). To further evaluate a role for DNA methylation in downregulating CEBPD expression, we treated a panel of AML cell lines including OCI-AML2, OCI-AML5 and KG1 with an optimized subtoxic dose (10% inhibitory dose, IC10; Supplementary Fig. 6E) of a clinically utilized hypomethylating agent, azacytidine, and assessed for changes in CEBPD gene expression. Consistent with the correlation analyses in the datasets, we found that azacytidine treatment was associated with upregulation of CEBPD expression (Supplementary Fig. 6F). These results further strengthen the potential role for DNA methylation in regulation of CEBPD genes expression.

In summary, our results suggest a tumor suppressor function for CEBPD in human AML cell lines. Our results offer further evidence to suggest that DNA methylation may regulate CEBPD gene expression in AML. Finally, our results suggest that CEBPD downregulation may contribute to the leukemic phenotype through activation of MAPK signaling. These results further improve our understanding of CEBPD function in AML biology.

## Service providers

Next generation sequencing services were provided by Novogene. Computational resources and technical support were provided by the Weill Cornell Medicine Applied Bioinformatics and the School of Medicine Research Computing at The University of Virginia. Flow cytometry services were provided by the flow cytometry core facility at the University of Virginia.

## Funding sources

NCI K08CA169055 (FGB), Pew-Stewart Scholar for Cancer Research supported by The Pew Charitable Trusts and the Alexander and Margaret Stewart Trust (FGB), NIH R35GM133712 (CZ), V-foundation for Cancer Research, the University of Virginia, and ASHAMFDP-20121 (FGB), UVA Comprehensive Cancer Center through the NCI Cancer Center Support Grant P30 CA44579 (FGB and SCP), Blood Cancer UK, Cancer Research UK and the UK National Institute of Health research (RD), Starr Cancer Consortium grant I4-A442 (AMM and RL), LLS SCOR 7006-13 (AMM) and The South Australian Cancer Research Biobank is supported by the Cancer Council SA Beat Cancer Project, Medvet Laboratories Pty Ltd and the Government of South Australia (RD).

We thank Drs. Feith, Tan, Goldfarb, Alagib, and Mohi, Samuel Haddox, and Emily Dennis from UVA for their technical advice and support.

## Affiliations

Tak Lee is currently affiliated with The Rockefeller University, Franck Rapaport is currently affiliated with Sanofi, Caroline Sheridan is currently affiliated with Immunai, and Michael Becker is currently affiliated with OhioHealth.

## Author contributions

Conceptualization: SCP and FEG-B

Data curation: SCP, ZW, YN, CZ and FEG-B

Formal analysis: SCP, ZW, YN, CZ and FEG-B

Funding acquisition: SCP and FEG-B

Investigation: SCP, ZW, CZ, and FEG-B

Methodology: SCP, ZW, YN, CZ and BP. Project administration: SCP and FEG-B

Resources: SCP, YN, CM, HF, CS, MWB, LB, MPC, RJD, RLL, AMM, CZ, and FEG-B

Software: SCP, ZW, ND, YN, CM, FA, and FEG-B

Supervision: SB and FEG-B

Validation: SCP, YN and FEG-B

Visualization: SCP and FEG-B

Writing – original draft: SCP and FEG-B

Writing – review & editing: all authors

## Data availability

RNA-sequencing from human AML patient specimens was obtained from dbGap accession number phs001027.v4.p1. Data generated from AML cell line OCI-AML5 will be deposited into GEO and can be made accessible to reviewers in a private Box folder during the review process.

