## Supplementary information for "CCAAT-enhancer binding protein delta functions as a tumor suppressor gene in acute myeloid leukemia"

**Supplementary information index**

**Methods**

**Chemical and Reagents**

A description of chemicals and reagents used in the study is provided Supplementary table 7.

**Cell lines**

HEK293FT cells were purchased from Thermo Fisher Scientific and used for all the lentiviral media preparations. AML cell lines SKNO-1, EOL-1, OCI-AML2, and OCI-AML5 were purchased from DSMZ. OCI-AML3 and Kasumi-1 cells were generous gifts from Dr. Loughran, University of Virginia Cancer Center. All cell lines were cultured per manufacturer's protocol (**S**upplementary table 7). All AML cell lines were authenticated using the human cell STR profiling service offered by ATCC.

**CEBPD knockdown and upregulation experiments**

We knocked down and upregulated CEBPD expression using a one-plasmid and two-plasmid CRISPR approach, respectively as described in our protocol (1), a modified protocol from a previous study (2). Briefly, to knockdown CEBPD expression, four independent CEBPD specific (sgCEBPD) and control (sgControl) sgRNAs (Supplementary table 8) (3, 4) were cloned separately into the BsmBI site of lentiCRISPRv2-GFP plasmid to prepare CEBPD knockdown and control constructs, respectively. The constructs were transfected individually with packaging plasmids psPAX2 and pMD2.G into HEK293FT cells to generate infectious lentiviral media. Equal volumes (250 µl) of the CEBPD specific or control lentiviral media were spinfected into equal number of OCI-AML5 cells (0.5 x 10^6^). After 3 days of spinfection, live GFP+ cells were sorted to establish polyclonal and monoclonal cell cultures. These cell cultures were used as stable CEBPD knockdown cells in downstream experiments after confirming CEBPD knockdown by WB. We validated CRISPR-mediated CEBPD knockdown results by shRNA-mediated CEBPD knockdown experiments. To knockdown CEBPD expression, we spinfected lentiviral media of three independent CEBPD specific (shCEBPD) and two independent control (shControl) shRNA plasmid constructs separately into OCI-AML2 and OCI-AML5 cells as described above for CRISPR experiments. Spinfected cell cultures were treated with puromycin (0.5 µg/ml) for seven days for selection of stably infected cells. Live spinfected cell cultures were used as stable CEBPD knockdown cells in downstream experiments after confirming CEBPD knockdown by WB.

To upregulate CEBPD expression, first, a stable OCI-AML2 or OCI-AML5 clone expressing transcriptional effectors MS2, p65, and HSF1 was generated by spinfecting lentiMPHv2 plasmid lentiviral media. Stably transduced cells were selected by treating with antibiotic hygromycin (250 µg/ml), and single cell sorted to generate lentiMPHv2 stable monoclonal cultures. Expression of transcriptional effectors in monoclonal cultures was assessed by RNA expression of one of the effectors MS2. One of lentiMPHv2 stable monoclonal cultures with high MS2 expression was selected for a second transduction in the next step. We used a sgRNA cloning plasmid lentiSAMv2 which expressed gene specific gRNA (Supplementary table 8) (2, 3, 5) and dCas9-VP64 to upregulate CEBPD expression. Three independent CEBPD activation (sgCEBPD) and control (sgControl) constructs were generated by cloning respective sgRNAs into the lentiSAMv2 plasmid. Individual constructs' lentiviral media were spinfected separately into the selected clonal lentiMPHv2 stable cells to upregulate CEBPD expression. Upregulation of CEBPD expression in the activation construct spinfected cells compared to controls was confirmed by qPCR and WB experiments.

**MTS Assay**

AML cells' growth rate was measured using CellTiter Non-radioactive Proliferation Assay (MTS) per manufacturer's protocol (Supplementary table 7). Briefly, equal numbers of viable control and CEBPD knockdown cells (2500 cells) were plated in 100 µL growth media into two separate 96 well plates, one plate for baseline and the other plate for endpoint reads. The baseline plate was read on the same plating day. Endpoint plate cells were cultured at 37°C in a CO_2_ incubator and read on day 5. Both baseline and endpoint plates were read colorimetrically at 570/670 nm using a BioTek Plate Reader. Baseline reads were subtracted from the endpoint reads to calculate net absorbance to estimate the change in cell growth rate. The student's t-test was used to assess statistical significance between control and CEBPD knockdown group means.

**Cell counting**

LUNA-FL Dual Fluorescence Cell Counter (Logos Biosystems) with AO/PI dye was used for routine cell culture counting and transduction experiments per manufacturer's recommendations (Supplementary table 7). For cell growth assays, cells were stained with 4',6-diamidino-2-phenylindole (DAPI) and viable cells were counted in 20 µl of cell suspension using Attune Nxt flow cytometer.

**Apoptosis**

Apoptosis rate in AML cells was measured by a flow cytometry-based method using APC-conjugated annexin V staining (Supplementary table 7). Equal number of CEBPD knockdown control and control cells were plated in a 12-well format plate at 0.5 x 10^6^ cells per ml density in their recommended growth media. After 5 days of culture, 0.2 million cells were aliquoted and washed, first with 1ml flow cytometry buffer and then with 0.5 ml annexin V binding buffer. Cells were resuspended in 100 µl annexin V binding buffer, and annexin V (1 µl) and DAPI (0.5 µl of 2.5 mg/ml) were added to the cell suspension. After 15 minutes of incubation at room temperature, an additional 200ul annexin V binding buffer was added, and APC signal was measured by flow cytometry. The student's t-test was used to assess statistical significance between control and CEBPD knockdown group means.

**EdU assay**

OCI-AML5 cells’ proliferation rate was assessed using Click-iT Plus EdU Alexa Fluor 647 Flow Cytometry Assay per manufacturer's protocol (Supplementary table 7). Briefly, equal numbers of control and CEBPD knockdown cells (0.5 x 10^6^) were plated in 1 mL media containing 10 µM EdU and incubated in a 5% CO_2_ incubator at 37°C. After 3 hours, cells were harvested and washed with 3 mL 1% BSA in PBS. Cells were fixed in 100 µL of Click-iT fixative for 15 minutes protected from light at room temperature. Fixed cells were washed with 3 mL 1% BSA in PBS and incubated with 100 µL of 1X Click-iT saponin-based permeabilization solution at room temperature. After 15 minutes, 500 µL Click-iT Plus reaction cocktail containing Alexa Fluor 647 picolyl azide was mixed with cells and incubated at room temperature protected from light. After 30 minutes, cells were washed with a 3 mL 1X Click-iT saponin-based permeabilization solution. Washed cells were resuspended in 300 µL of 1X Click-iT saponin-based permeabilization solution and analyzed for EdU incorporation. An Attune NxT Flow Cytometer was used for data collection, and analysis was performed using FCS Express 7 software. EdU untreated cells served as control to gate EdU positive cells. The student's t-test was used to assess statistical significance between control and CEBPD knockdown group means.

**Colony-forming unit assay**

Self-renewal capacity of control and CEBPD knockdown OCI-AML5 cells was compared using a colony-forming unit (CFU) assay. Equal number of control and CEBPD knockdown cells (~100 cells\well) were plated in duplicates in a 6-well plate format in 1.1 ml of methylcellulose-based MethoCult Classic medium (Supplementary table 7). After 7 days of culture, colonies were counted manually under an inverted microscope. Cells were harvested, and equal number of control and CEBPD knockdown cells (~100 cells\well) were re-plated in a 6-well plate format in 1.1 ml of methylcellulose-based MethoCult Classic medium. Re-plating was repeated up to 6 times to assess the self-renewal capacity of CEBPD knockdown compared to control cells. The student's t-test was used to assess statistical significance between control and CEBPD knockdown group means.

**Quantitative reverse transcription chain reaction (RT-qPCR)**

RT-qPCR was used to assess changes in genes' mRNA expression. Total RNA was purified from 1x10^6^ cells using Trizol method as described previously (1).  Purified RNA (~1 µg) was converted to cDNA using the M-MuLV reverse transcriptase enzyme in an optimized reaction mixture. Genes' expression was measured in an optimized reaction mixture containing Q5 Hot Start Master Mix, EvaGreen, gene specific primers (Supplementary table 7), and 40 µg of the converted cDNA (~40 µg RNA) using StepOne RT-qPCR machine. The student's t-test was used to assess statistical significance between control and CEBPD knockdown group means.

**Western blot**

Western blot (WB) analysis was performed to assess protein expression and phosphorylation levels using enhanced chemiluminescence (ECL) or Licor method. Briefly, AML cells (1x10^6^) were lysed in RIPA buffer (100 µL) containing protease inhibitor cocktails. Lysed cells were centrifuged at 4°C at 14000 rpm for 10 minutes. Supernatant was transferred to an Eppendorf tube and used as cell lysate. Equal amounts of control and experimental cells' lysates (30 µg) were resolved in 4% stacking and 15% resolving Tris-glycine polyacrylamide gel (50V for 30 minutes and 100V until dye reaches the bottom of gel at room temperature) and transferred to a 0.2 µM PVDF membrane (1X Tris/Glycine buffer with 200 ml methanol/Liter). PVDF membrane was blocked with 5% nonfat dry milk in PBS (ECL method) or Licor buffer (Licor method) for 1 hour, incubated overnight at 4°C with optimized concentration of a primary antibody (Supplementary table 7). On the next day, membranes were washed and incubated with a secondary antibody (Supplementary table 7) in 5% non-fat dry milk in PBS (ECL method) or Licor buffer (Licor method) and developed with an enhanced chemiluminescence substrate or Licor imaging system.

**Azacytidine treatment**

Azacytidine kill curves were generated by treating parent AML cell lines (0.5x10^6^/mL) with a range of azacytidine concentration (0-6400 nM) every day for 5 days and measuring cells' viability by MTS assay. The curves were used to determine cell line specific sub-toxic concentration of azacytidine which reduced cell line's cell viability by 10% (IC_10_). To assess the effect of azacytidine treatment on CEBPD expression, AML cell lines were treated with sub-toxic doses of (IC_10_) for 5 consecutive days. After treatment, CEBPD mRNA expression was assessed by RT-qPCR. The student's t-test was used to assess statistical significance between control and CEBPD knockdown group means.

**Bulk RNA-Seq analysis**

PAML patients' bulk RNA sequencing raw data was downloaded from dbGaP (accession phs001027.v4.p1). It comprised of two cohorts (I: n = 29 and II: n = 30). Each cohort comprised of matched diagnosis-relapse pairs along with corresponding healthy specimens from CD34+ normal bone marrow cells (Supplementary table 1). Library preparation and sequencing were previously described (6, 7). Sequencing reads were aligned to the human genome assembly GRCH37/hg19 using STAR v2.4.0f1, annotated using subread v1.4.5-p1 featureCounts and read counts were normalized to Transcripts Per Million reads (TPM). Differentially expressed genes were calculated using DESeq2 by comparing each diagnosis or relapse sample and including the batch-matched healthy samples in the model matrix for normal variance estimation. Levels of differentially expressed genes were batch corrected using limma package to estimate combined changes in genes’ expression upon disease relapse in all the samples from both the cohorts. Significantly differentially expressed genes (DEGs) were selected at thresholds of adjusted *p*-value (q-value) <0.01 and log2 fold change > |1.2|. AML cell lines' mRNA sequencing was performed using ABclonal Fast RNA-seq Lib Prep Kit V2 and sequenced on a NovaSeq X Plus machine using pared end 150 base pair sequencing at Novogene. Sequencing reads were aligned to the human genome assembly GRCH38/hg38 using hisat2 (8) and annotated using htseq-count (9). Differential gene expression analyses between CEBPD knockdown and control samples were performed using DESeq2. Significantly differentially expressed genes were selected at thresholds of adjusted *p*-value < 0.05.

**Gene set enrichment analysis**

Gene set enrichment analysis was performed by using a fast gene set enrichment analysis (fgsea) package in R (10). The genes were first pre-ranked by the log2 fold changes and then assessed for enrichment using the human h.all.v7.5.symbols.gmt gene set. Treatment sensitivity gene sets were differentially expressed gene lists determined between resistant and sensitive patient specimen groups as the baseline identified in Beat AML's functional treatment response data (11, 12). These Beat AML gene sets were kindly provided by Dr. Jeffrey Tyner. Molecular pathways with adjusted p-value < 0.05 were considered significantly enriched upon CEBPD knockdown.

**BART**

We used log2 fold change values of all the genes that were differentially expressed (log2FoldChange > |1| & q-value < 0.01) in at least 5 patients to identify 6 gene clusters in each cohort, first via up or downregulation from normal to diagnosis, and then second into upregulation, downregulation, or diverse expression changes from diagnosis to relapse. The consistently downregulated and the consistently upregulated gene sets largely separated the patients into two groups for each cohort.  The consistently downregulated gene set included 1085 genes for cohort I and 817 genes for cohort II, respectively, with 519 genes shared. We used BART (version 2.1) (13) to predict transcriptional regulators that function to regulate each set of differentially expressed genes. CEBPD was identified as one of the potential regulators for the 519 consistently downregulated genes shared between the two cohorts (p-value < 0.05).

**Publicly available datasets**

publicly available datasets Beat AML (11, 12), TCGA (14), TARGET (15), and Hackl et al (16) were used as validation datasets. The datasets were downloaded from the publicly available data portals GDC (TCGA & TARGET), github repository (Beat AML, <https://biodev.github.io/BeatAML2/)>, and GSE66525 (16).

**Methylation analysis**

Study specimens' Enhance Reduced Representation Bisulfite Sequencing (ERRBS) data was downloaded from dbGaP (accession phs001027.v4.p1). Library preparation, bisulfite conversion, sequencing, alignment, and methylation calls were described previously (17). The mean of methylation scores of non-converted cytosines between -250bp to +500bp from the TSS in each specimen was used in correlation analyses to assess association with CEBPD gene expression for TCGA and Beat AML datasets.  The mean of methylation scores of non-converted cytosines for the gene was used in correlation analyses to assess association with CEBPD gene expression for TRAGET dataset. Publicly available datasets were downloaded from publicly available data portals. A robust linear model (RLM) was employed to calculate the line of best fit in the correlation plot. Spearman test was used to assess statistical significance of the association.

**Copy number alteration and mutation**

CEBPD copy number alteration and mutation analyses were performed using cBioPortal web application (https://www.cbioportal.org/). Acute Myeloid Leukemia (TCGA, NEJM 2013, Acute Myeloid Leukemia (OHSU, Nature 2018), and Acute Myeloid. Leukemia (TARGET, GDC) datasets were used for TCGA, Beat AML, and TARGET dataset analyses, respectively. Plots for the PAML dataset (18) were generated manually using Microsoft PowerPoint.

**Survival analyses**

Patients’ overall survival analyses were performed using the Kaplan-Meir method. Patients who were treated with standard AML chemotherapy 7+3 regime plus additional agents were included in the analysis. The top and bottom quartiles of patients based on CEBPD expression were classified as CEBPD-High and CEBPD-Low respectively. The log-rank test was used to assess statistical significance of association.

**Statistical analyses**

The statistical significance of the results was assessed using the Graph Pad or the R software. Wilcox tests, t-tests, and spearman tests were used for boxplots, bar graphs, and correlation analysis comparisons, respectively. Statistical thresholds: *p-value < 0.05; **p-value < 0.001; ***p-value < 0.0001.

**Supplementary figures and legends**

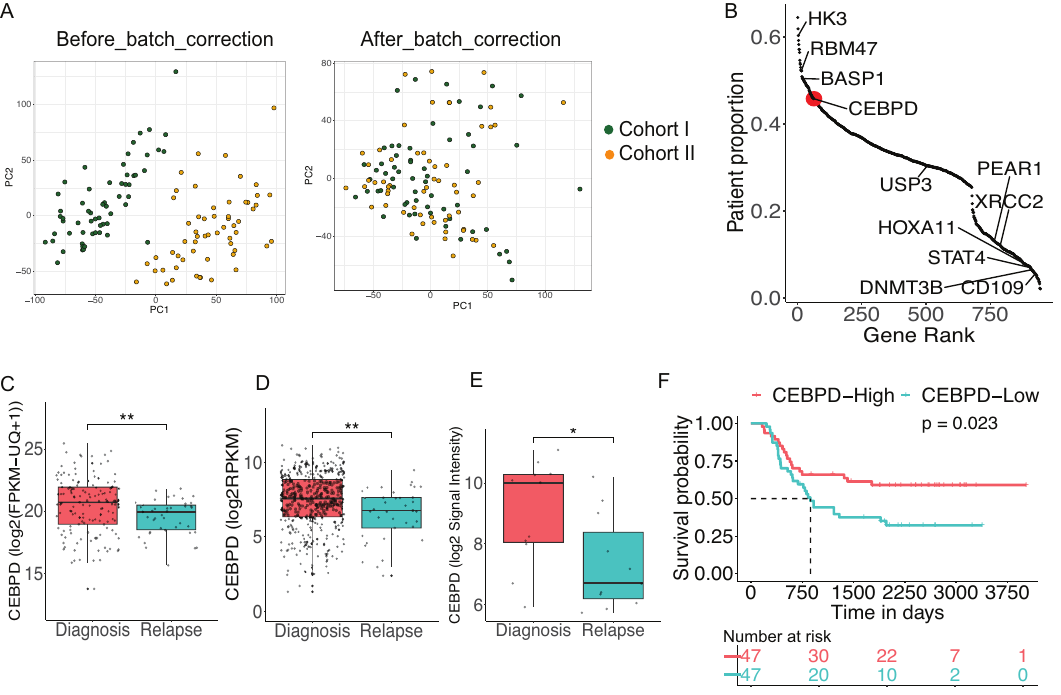

**Supplementary figure 1. Loss of CEBPD expression is associated with relapse and poor survival of AML patients.** (**A**) Results of principal component analysis before and after application of batch correction for the two cohorts (I: green & II: yellow) of the study dataset. (**B**) Proportion of patients exhibiting significantly differentially expressed genes in the study dataset. Gene Rank is an arbitrary rank of the gene. CEBPD is shown with a red dot. (**C-E**) A comparison of CEBPD mRNA expression between diagnosis and relapse patients in TARGET (C), Beat AML (D), and Hackl et al (E) datasets. Wilcox test was used to determine statistical significance between the two groups. (**F**) Kaplan Meir survival analysis results for association between CEBPD expression and overall survival in the TARGET AML dataset. Log-rank test was used to determine statistical significance of association.

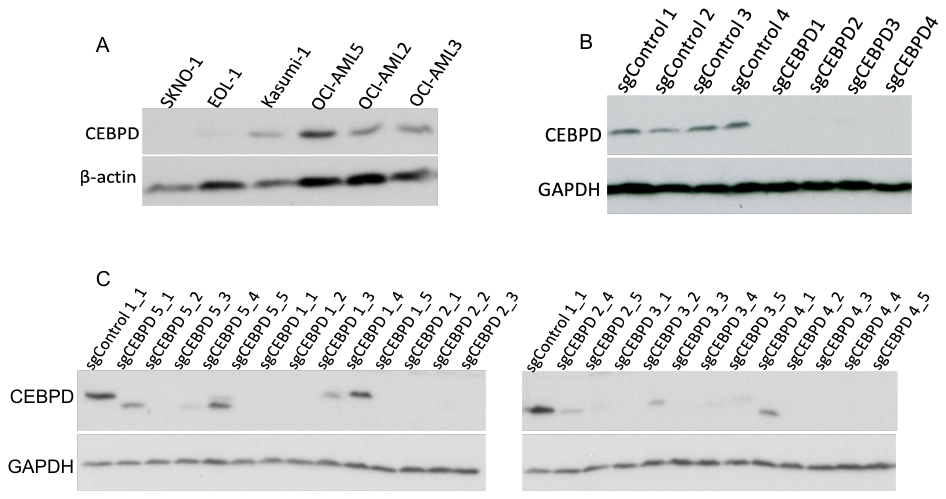

**Supplementary figure 2. CRISPR-mediated approach knocked down CEBPD expression efficiently in OCI-AML5 cells.** (**A**) Western blot (WB) results showing CEBPD expression levels in AML cell lines. β-actin served as a total protein loading control. (**B-C**) WB results showing CEBPD expression levels in CEBPD knockdown stable polyclonal (B) and clonal (C) OCI-AML5 cells. CEBPD expression levels in polyclonal cells transduced with control (sgControl) and CEBPD targeting sgRNA plasmid constructs (sgCEBPD) are shown for sgControl 1 to 4 and sgCEBPD 1 to 4, respectively. CEBPD expression in clonal CEBPD knockdown cells is shown for sgCEBPD 1 to 5. Numbers 1 to 5 after the underscore represent clone numbers for the individual CEBPD targeting sgRNA plasmid constructs. Clonal sgControl 1_1 was used as control. GAPDH served as total protein loading control.

**Supplementary figure 3. Knockdown of CEBPD expression enhanced AML cell growth, while overexpression induced expression of myeloid differentiation marker.** (**A**) WB results showing CEBPD expression in CEBPD knockdown (shCEBPD 1 to 3) compared to control (shControl 1 to 2) OCI-AML2 cells. (**B**) Flow cytometry-based live cell counts in CEBPD knockdown compared to control OCI-AML2 cells. (**C**) WB results showing levels of CEBPD expression in CEBPD knockdown compared to control OCI-AML5 cells. (**D**) Flow cytometry-based live cell counts CEBPD knockdown compared to control OCI-AML5 cells. (**E**) Representative picture of 6-well format plate well plated with CRISPR-mediated CEBPD knockdown (sgCEBPD) compared to control (sgControl) cells at repeated plating 6. (**F**) Representative Annexin V apoptosis assay cell density plots in CEBPD knockdown compared to control cells. OCI-AML5 cells which were not stained with Annexin V-APC served as gating control. (**G-H**) Results of qPCR analyses showing levels of CEBPD (G) and CD14 (H) mRNA expression in CRISPR mediated CEBPD activated (sgCEBPD) compared to control (sgControl) OCI-AML5 cells. (**I**) WB results showing levels of CEBPD expression in sgCEBPD activated and sgControl OCI-AML5 cells. (**J**) Results of qPCR analyses showing levels of CD14 mRNA expression in sgCEBPD activated compared to sgControl OCI-AML5 cells. The student's t-test was used to determine the statistical significance between the two groups.

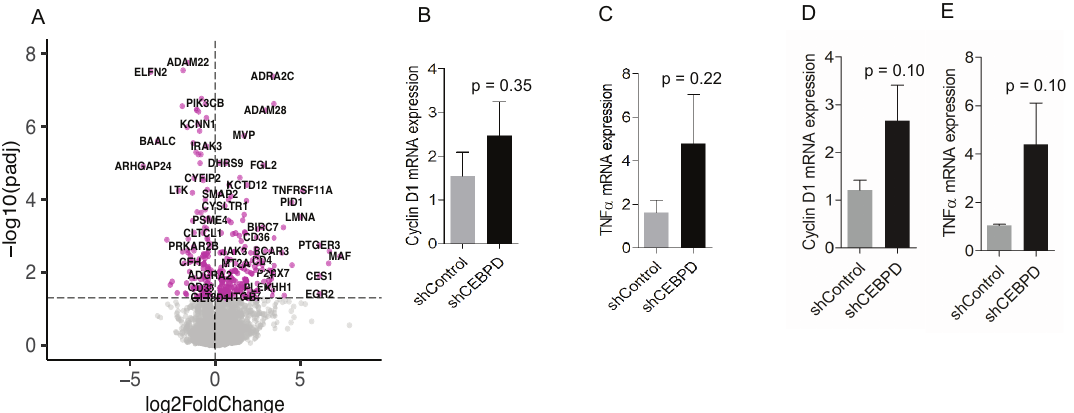

**Supplementary figure 4. Knockdown of CEBPD expression activated MAPK signaling transducers cyclin D1 and TNFα.** (**A**) Volcano plot showing differentially expressed genes upon CRISPR-mediated CEBPD knockdown in clonal OCI-AML5 cells. The differential gene analysis was performed between control (sgControl 1 to 4) and CEBPD knockdown (sgCEBPD 1 to 4) cells. Dotted horizontal line represents statistical threshold padj < 0.05. (**B-E**) Results of qPCR analyses showing cyclin D1 and TNFα mRNA expression in OCI-AML2 (B-C) and OCI-AML5 (D-E) cells in shRNA-mediated CEBPD knockdown (shCEBPD 1 to 3) compared to control (sgControl 1 to 2) cells. Add statistics to figure B-E. The student's t-test was used to determine the statistical significance between the two groups.

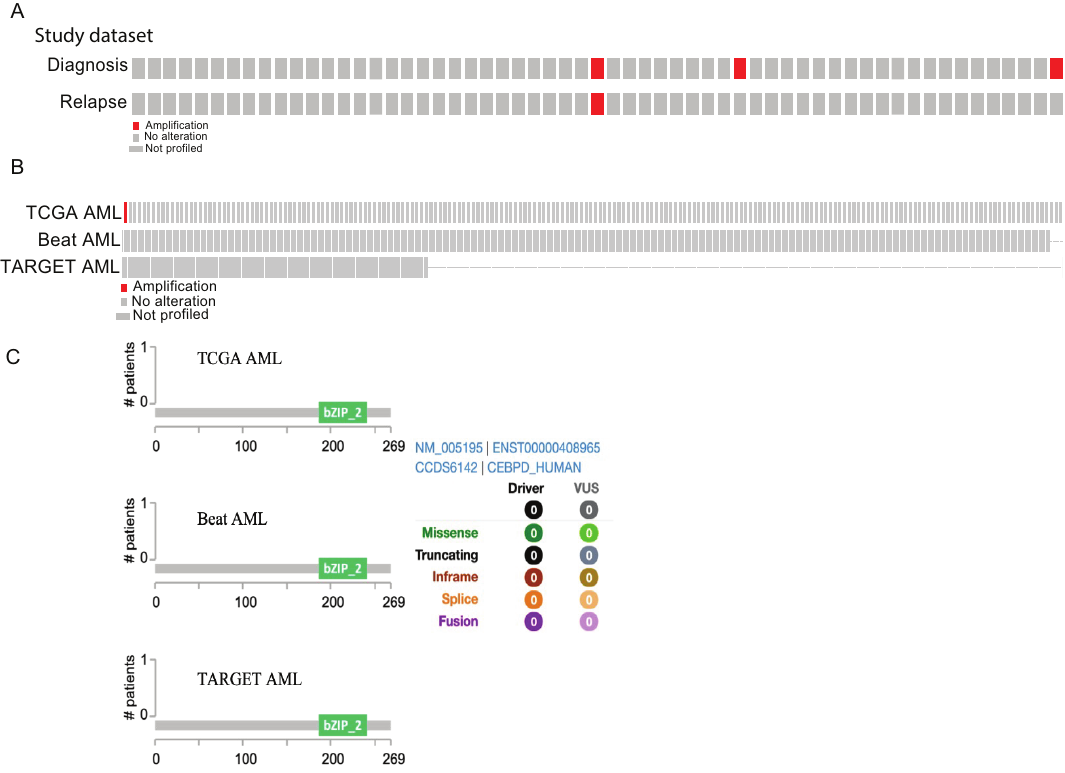

**Supplementary figure 5. Downregulation of CEBPD expression is not associated with genetic alterations in AML patients.** (**A-B**) Oncoprint plot from CEBPD copy number alternation analyses (CNA) performed on the study (A) and publicly available datasets (B). The plot for the study and publicly available datasets were generated using Microsoft Power Point and cBioportal web application, respectively. Vertical red and grey bars represent amplification and no alteration, respectively. Horizontal grey bars represent specimens that were not processed for CNA. (**C**) Lollipop plots from CEBPD mutation analyses performed on publicly available datasets using cBioportal web application. Types of mutations are color coded.

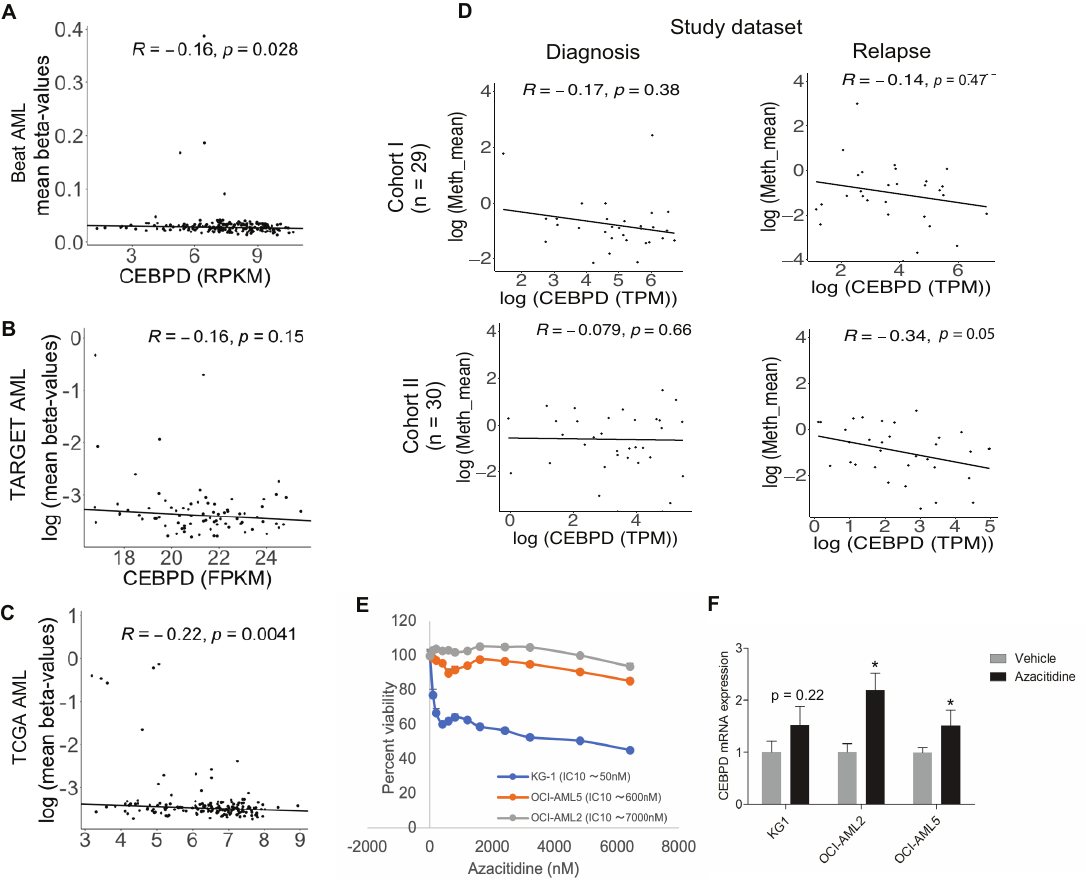

**Supplementary figure 6. Loss of CEBPD expression is associated with a trend of promoter methylation.** (**A-D**) Results of correlation analyses between CEBPD mRNA expression and promoter (-250bp to +500bp from the TSS) methylation in the publicly available (A-C) and the study datasets (D). Spearman's test was used to determine statistical significance of the association. (**E**) Azacitidine kill curves of KG1, OCI-AML2, and OCI-AML5 cells. IC10 represents azacitidine dose that inhibited growth of cells by 10%. (**F)** Results of qCR analyses showing CEBPD mRNA expression in azacitidine compared to vehicle treated AML cell lines. GAPDH expression served as reference for the read normalization. The student's t-test was used to determine statistical significance between the two groups.

**Supplementary table legends**

**Supplementary table 1. Study dataset specimens' description.** Patient ID = patient identifying term. Disease stage =  patient’s disease stage identifier at the time of specimen collection. Cohort annotation = patient’s cohort number identifier.

**Supplementary table 2. Results (log2FC) of DESeq2 analysis on relapse compared to diagnosis specimens in the study dataset.** Column A = gene identifier. Column B-BH = patient identifier. Column Upregulated_Number = number of patients with log2FC > 1.2. Column Downregulated_Number = number of patients with log2FC < -1.2. Column Upregulated_Proportion = Upregulated_Number/Total number of patients. Column Downregulated_Proportion = Downregulated_Number/Total number of patients.

**Supplementary table 3. Results of DESeq2 analysis on diagnosis compared to normal bone marrow control specimens in the study dataset.** Gene ID = gene identifier. Patient ID = patient identifier. DvsN_log2FC = Diagnosis vs Relapse specimen log2FC, DvsN_qvalue = Diagnosis vs Relapse specimen qvalue.

**Supplementary table 4. Results of DESeq2 analysis on relapse compared to diagnosis specimens in the study dataset.** Gene ID = gene identifier. Patient ID = patient identifier. RvsD_log2FC = Diagnosis vs Relapse specimen log2FC, RvsD_qvalue = Diagnosis vs Relapse specimen qvalue.

**Supplementary table 5. Results of BART analysis.** TR = transcriptional regulator.  statistic, pvalue, zscore, max-auc, re_rank, irwin_hall_pvalue are the statistical parameter outputs from BART analysis.

**Supplementary table 6. Results of DESeq2 analysis on CEBPD knockdown (sgCEBPD 1 to 4) compared to control (sgControl 1 to 4) clonal OCI-AML5 cells.**  gene = gene identifier. baseMean, log2FoldChange, lfcSE, stat, pvalue, and padj are statistical parameter outputs from the DESeq2 analysis.

**Supplementary table 7. Description of the chemical and reagent used in the study.**

**Supplementary table 8. Description of sgRNA sequences used in the study.**
